# Comparison of African, Asian, and American Zika Viruses in Swiss Webster mice: Virulence, neutralizing antibodies, and serotypes

**DOI:** 10.1101/075747

**Authors:** Charles B. Stauft, Oleksandr Gorbatsevych, Jeronimo Cello, Eckard Wimmer, Bruce Futcher

**Affiliations:** Dept. of Molecular Genetics and Microbiology, Stony Brook University, 11794-5222; Codagenix Inc., Stony Brook, NY 11794

## Abstract

The sequence of Zika virus has evolved as it has spread out of Africa and into the Americas. It is unclear whether American strains of the virus define a new serotype. Here, we have tested the virulence and immunogenicity of three wild-type ZIKV strains in neonatal Swiss Webster mice. We found that all three ZIKV strains (African MR766, 1947; Asian FSS13025, 2010; and American, PRVABC59, 2015) are capable of killing neonatal mice after intracranial injection. Intraperitoneal injection with these viruses did not kill, but produced neutralizing antibodies as measured by a PRNT_50_ assay. Sera from mice infected with each virus were tested for neutralizing activity against the infecting virus and also the other two viruses by a PRNT_50_ assay. In general, the antibodies induced by each virus were good at neutralizing that virus (the homologous virus), but somewhat poorer at neutralizing the other two viruses (heterologous viruses). Antibodies induced by the African strain MR766 were about 4-fold worse at neutralizing the American strain PRVABC59 than the homologous strain, while antibodies induced by the American strain were about 10-fold worse at neutralizing the African strain than the homologous strain. Because the antibodies are cross-neutralizing at some level, the viruses do not form separate serotypes. Nevertheless, these results raise concern that the immunity conferred by the African virus may protect only relatively poorly against the new American strains. This has implications for the possible spread of the American ZIKV strains to Africa and Southeast Asia, and also for the development of vaccines.

## Introduction

Zika virus has migrated out of Africa where it was discovered in 1947[1] into Southeast Asia[2–4], into Micronesia [5, 6] and French Polynesia [7, 8] and most recently into South and Central America [9–13]. Since 2015, mosquito-borne Zika virus transmission has been reported in 64 countries and territories, with 50 countries and territories reporting their first ever Zika virus outbreaks (WHO). Alarmingly, Zika virus infection causes birth defects such as microcephaly in infants [14–16] and Guillain-Barre syndrome in adults [17–19]. Zika virus has also reached the United States, with 4,729 locally acquired cases in US Territories and 1,657 travel-associated cases reported within the United States itself[20]. Zika virus infection within the United States and its territories are already associated with 22 cases of Guillain-Barre syndrome. 885 infections are of pregnant women [21] whose children are now at risk for developing birth defects.

Zika virus was first discovered in Uganda in 1947. Subsequence serological studies suggested that, at least in some regions of Africa, a very large proportion of the adult population is seropositive for Zika virus, and so is presumably immune to new infections. However, as Zika virus has spread across the globe, its primary genome sequence has changed. Because of the significant amino acid changes from the ancestral MR766 to the new American strains, a question arises as to whether the American strains of virus form a novel serotype. That is, will individuals with immunity against the ancestral African virus also have cross-immunity against the new American strains? If not, there is a concern that the novel strains may cause a wave of infection across Africa and Southeast Asia, notwithstanding the fact that older forms of Zika virus are already endemic there. Furthermore, there is an issue of whether a vaccine for Zika virus should be based on the MR766 amino acid sequence.

Zika virus, like dengue virus, infects primates[1, 22]. However, neonatal wild-type mice are also sensitive to Zika [23] and dengue [24] viruses. Previously, we have used a proxy challenge model using immune competent mice to gauge the effectiveness of dengue virus vaccine candidates to protect neonatal mice against lethal challenge [25]. We feel that this model will be equally applicable to research with ZIKV, a closely related flavivirus. Here, we infect neonatal mice with Zika. After intracranial injection, these infections are lethal, while after intraperitoneal injection, infections induce neutralizing antibodies. We compare the African, Asian, and American viruses for virulence in this model, and ask whether the antibodies induced by one strain differ in potency against the other strains.

## Materials/Methods

### Cells and Viruses

Vero cells were grown in Dulbecco's Modified Eagle's Medium (DMEM) supplemented with 10% bovine calf serum (BCS; Gemclone) and Penicillin/Streptomycin (CellGro).

ZIKV strain MR766-SM150 (LC002520) and FSS13025 (JN860855) were acquired from Dr. Robert Tesh (UTMB, Galveston TX). ZIKV strain PRVABC59 (KU501215) was provided by BEI Resources (NIAID, NIH: Zika Virus, PRV-ABC59, NR-50240). MR766[1] was isolated in 1947 from a primate in Uganda, FSS13025[25] was isolated in Cambodia from a human patient in 2010, and PRVABC59[26, 27] is a recent isolate (2015) from a human patient in Puerto Rico. Before infection in animals, stocks of each virus were generated by infecting confluent Vero cells at an MOI of 0.1 and harvesting cell culture supernatant at 3 days post infection.

### Mouse Infection

To prepare immune serum, African (MR766 P3) or Asian (FSS13025) or American (PRVABC59) Zika viruses were used to infect 4 week old Swiss Webster (Charles River) mice at a dose of 1 x 10^6^ PFU delivered intraperitoneally in a volume of 100 μL DMEM. Six mice were infected with each virus, but one of the mice infected with FSS13025 failed to develop any antibody titer against any virus (perhaps due to an injection failure), and this serum was not included in analysis. At 28 days' post-infection, mice were euthanized and terminally bled through cardiac puncture. Whole blood was allowed to clot for 30 minutes at room temperature, centrifuged at 10,000 rpm for 15 minutes, serum collected and frozen at −80°C.

To determine LD_50_ in newborn Swiss Webster mice, litters (n=7–12) were intracranially injected with 20 μL of ZIKV MR766, FSS13025, or PRVABC59 diluted in Dulbecco's Modified Eagle's Medium (DMEM). Mice were checked daily for morbidity and mortality for 21 days post infection. All animal experiments were conducted under approval by the Institutional Animal Care and Use Committee at Stony Brook University.

### PRNT_50_Assay

Standard working solutions of MR766 (Monkey / Uganda / 1947), FSS13025 (Human / Cambodia / 2010), and PRVABC59 (Human / Puerto Rico / 2015) were prepared at a concentration of 500 PFU/mL. Mouse serum was heat treated for 30 minutes at 56°C and then mixed with 100 PFU of each virus separately with 4-fold serum dilutions of 1:10, 1:40, 1:160, and 1:640 diluted in Vero infection medium (5% BCS DMEM). Each serum/virus mixture was incubated at 37°C for 2 hours and then added to confluent Vero cell monolayers in a 12-well format with a volume of 250 μL. Plates were rocked at room temperature for 30 minutes, then 250 μL of additional Vero infection medium was added and the plates incubated at 37°C, 5% CO_2_ for 1 hour. Medium was then aspirated from each plate, the wells were washed once with sterile DPBS, and 1 mL of 0.6% Tragacanth Gum, 5% BCS DMEM was added to each well. The plates were incubated at 37°C, 5% CO_2_ for 4 days before being stained with crystal violet and read. The PRNT_50_ titer was determined by taking the reciprocal of the highest dilution that reduced the plaque count by 50% or more. PRNT_50_ values were log_2_ transformed and compared using a paired t-test for MR766 and PRVABC59 infected mice, for which the virus-to-virus differences in PRNT_50_ values were distributed normally. There was not a normal distribution for the mice infected with FSS13025, so these results were not analyzed with the paired t-test.

## Results and Discussion

Alignment of the sequence from the African strain MR766 (Uganda, 1947) with the American strain PRVABC59 (Puerto Rico, 2015) shows about 10% nucleotide change. A pairwise alignment of the polyprotein amino acid sequences for MR766 and either PRVABC59, or FSS13025 reveal a similar mutation profile where both the American and Asian strains contain about 125 amino acid differences from the African strain polyprotein (Fig. 1 B). Interestingly, both PRVABC59 and FSS13025 contain a four amino acid insertion in the envelope E protein that is responsible for host cell receptor binding (Fig. 1 B in red). FSS13025 is closely related to PRVABC59, containing only one amino-acid substitution in the E protein. Swiss Webster neonatal mice are highly susceptible to DENV[28] and ZIKV[23] when delivered intracranially. In preparation for future vaccine studies, we established the virulence of wild-type Zika viruses using this model. Individual litters were infected and mice were weighed and checked daily for signs of infection. The most commonly observed sign of infection was piloerection and the Swiss Webster neonates were susceptible to infection by all three strains of ZIKV (Fig. 2). Our strain of MR766 was passaged 150 times in suckling Swiss Webster mice [23] and was very lethal in this model with an LD_50_ of 1 PFU, possibly because of this passaging. FSS13025 and PRVABC59 were also lethal in newborn mice, with LD_50_ values of 250 and 2.7, respectively (Table 1) without recorded passaging in newborn mice. Thus, PRVABC59 is almost as virulent in mice as MR766, despite the fact that the latter has been mouse-adapted by passaging, and the former has not. Mean time to death (MTD) was also calculated for the highest dose administered in each group (Table 1). Similar in pattern to the LD_50_ results, MR766 infected mice succumbed more quickly to infection (MTD=6.91 days) compared to FSS13025 (MTD= 13.0 days) or PRVABC59 (MTD= 11.4 days).

**Figure 1.**
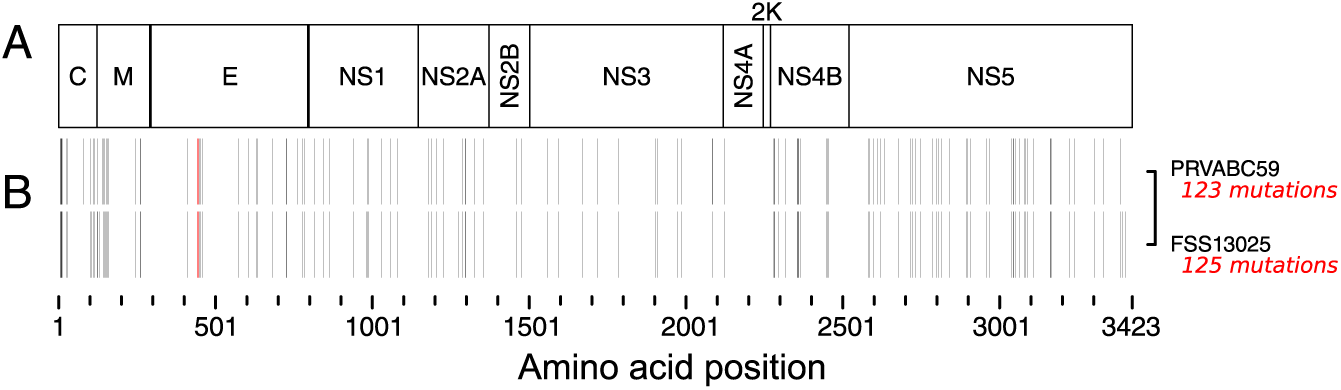
Mutation profiles of FSS13025 and PRVABC59 when aligned to MR766. Single amino acid differences are denoted by black and red bars relative to the polyprotein sequence of MR766 (B). The four red bars highlight a four amino acid insertion in FSS13025 and PRVABC59. Mutations map to the polyprotein products outlined in (A).

**Figure 2.**
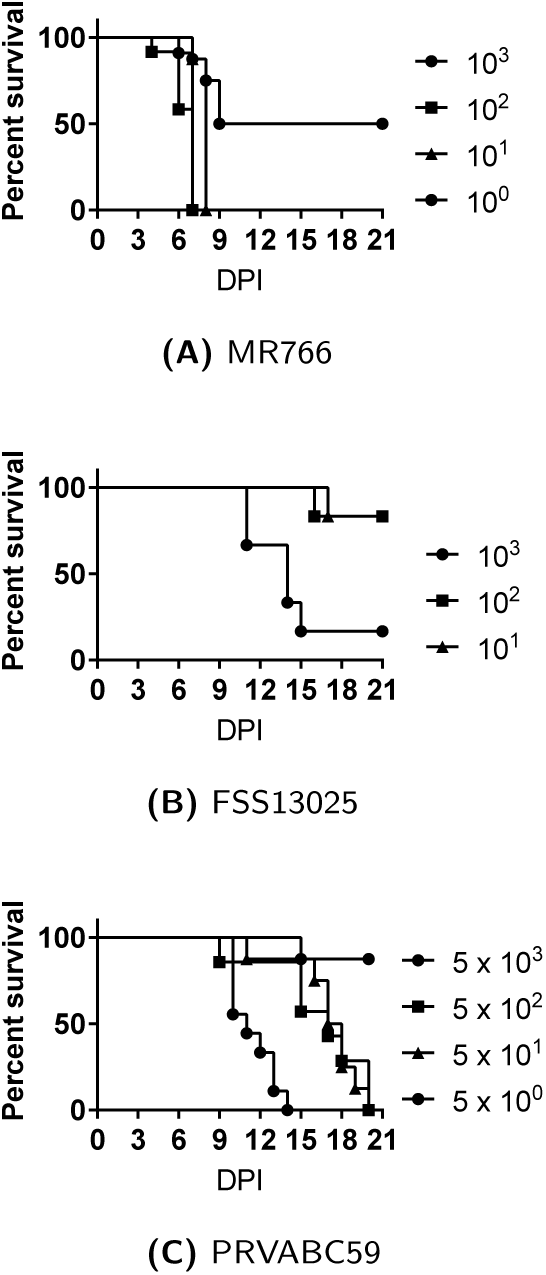
Neonatal Swiss Webster mice were infected with MR766 (A), FSS13025 (B) or PRVABC59 (C) injected intracranially.

**Table 1.** Neonatal, 1–2 day old, Swiss Webster mice were infected intracranially with serial dilutions of MR766, FSS13025, or PRVABC59 and mortality assessed using Kaplan-Meier curves. LD_50_ and mean time to death (MTD) values were calculated for the highest dose used in the infections.

| Virus | LD<sub>50</sub> | MTD |
| --- | --- | --- |
| MR766<sup>a</sup> | 1 | 6.91 |
| FSS13025<sup>b</sup> | 250 | 13 |
| PRVABC59<sup>c</sup> | 2.7 | 11.4 |

<sup>a</sup> $1 \times 10^3$ PFU dose
<sup>b</sup> $1 \times 10^3$ PFU dose
<sup>c</sup> $5 \times 10^3$ PFU dose

Six mice were infected intraperitoneally with the African and American viruses, and five mice with the Asian virus, and immune sera were harvested. All sera were tested by PRNT_50_ assay for neutralizing antibody titers against all three viruses (Fig. 3). Titers from mice injected with the African strain were 4-fold higher against the homologous African strain than against the heterologous Asian or American strains (Fig. 3A, Tables S1-S3). These differences were statistically significant (p = 0.042 for each comparison) by a paired t-test. Conversely, titers from mice injected with the American strain were 10-fold higher against the homologous American strain than against the heterologous African strain (Fig. 3C, Tables S1–S3), and about 1.6-fold higher than against the Asian strain (Fig. 3C, Tables S1–S3). The 10-fold difference between the American and African strain titers was highly significant (p = 0.014) by a paired t-test. Results with mice injected with the Asian strain were intermediate, but appeared qualitatively more similar to results with the American strain than the African strain (Fig. 3B). Because titer differences with the Asian virus were not normally distributed, significance could not be evaluated by the paired t-test.

**Figure 3.**
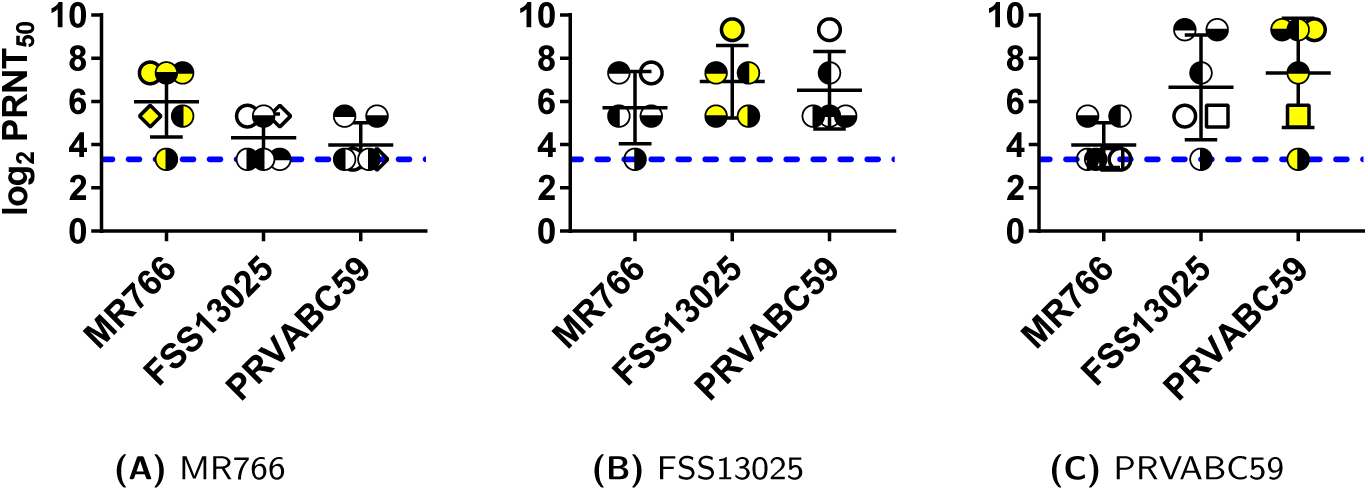
4-week old, Swiss Webster mice were infected with 1 × 10^6^ PFU of either the MR766, FSS13025, or PRVABC59 strains of ZIKV. Serum was collected from each mouse at 28 days post infection and tested by PRNT_50_ against homologous and heterologous strains of ZIKV. In each panel (A, B, or C), results from serum from an individual mouse are marked with a different symbol; i.e., an open circle with no fill represents a single mouse in that panel. Yellow shading in symbols indicates the homologous virus.

These results show that there is significant strain specificity in the neutralizing antibodies induced by African and American strains: Antibodies raised against the African strain are more potent (˜4-fold) against the African than the Asian or American strains, while antibodies raised against the American strain are more potent (˜10-fold) against the American strain than the African strain (Tables S1-S3). This is consistent with the sequence divergence between these strains. The fact that these strains can be distinguished serologically needs to be taken into account in vaccine design: a vaccine against the American strains is more likely to be effective if it is based on an American strain than if it is based on an African strain.

Our serological results so far are based on the in vitro PRNT_50_ assay. An obvious next step would be to look at in vivo protection. Neonatal mice that have acquired antibodies induced by one virus (either by direct injection of serum, or maternal antibodies)24 could be challenged with the homologous and heterologous viruses.

A recent paper correlated neutralizing antibodies[27] against Zika virus (specifically anti E glycoprotein antibodies) with protection against challenge. Using tests for neutralizing antibodies in infected mice is therefore essential to testing vaccine candidates. Larocca et al[27] noted that a microneutralization titer (essentially a PRNT_100_ value) of 10 was sufficient to protect mice against challenge. The limit of detection (blue line) for that the PRNT_50_ titers we observed, while low, are still sufficient to protect mice against heterologous challenge with other strains of ZIKV.

Two recent papers have reached conclusions somewhat different from ours. First, Aliota et al. [29] infected rhesus macaques with the African Zika strain MR766, and challenged these macaques 70 days later with a French Polynesian strain. The macaques were fully protected against the second infection, showing no viremia. Aliota et al. concluded that ``Immunogen selection is unlikely to adversely affect the breadth of vaccine protection'' and ``ZIKV strain selection is unlikely to compromise vaccine effectiveness''. However, protection after 70 days (while certainly a good thing!) is not a quantitative measure of vaccine effectiveness. At 70 days after the primary infection, antibody titers are likely very high, and likely to afford protection even if those antibodies are not perfectly matched against the new antigens. We found a 4 to 10-fold strain-dependent difference in the neutralizing activity of our sera, which does not immediately conflict with Aliota et al as these titers may still be protective in vivo[27]. Regardless of the difference we feel that for an optimal vaccine, immunogen selection is likely important.

Second, Dowd et al.[30] did mouse experiments broadly similar to ours, with several differences in technical detail, but unlike us did not find statistically-significant differences in cross neutralization. Possibly relevant technical differences are that we compared a recent American virus, PRVABC59, to the African virus MR766, while Dowd et al. compared an Asian virus from 2013 (H/FP/2013). In addition, we infected juvenile wild-type mice to obtain sera, while Dowd et al. infected mutant irf3/irf3-/-mice. However, it is not clear that either of these technical differences should matter. We saw moderate differences in PRNT_50_ (i.e., neutralizing antibody titer) between viruses of four-fold (when MR766 was the infecting virus) to ten-fold (when PRVABC59 was the infecting virus). We judged these differences to be statistically significant using a paired t-test, which is an appropriate and relatively powerful test under these circumstances. In contrast, Dowd et al. saw essentially no differences when MR766 was the infecting virus, and differences of perhaps two-fold when the Asian virus H/PF/2013 was the infecting virus (Fig. 4A of Dowd et al.), and they judged these differences to be not statistically significant using ANOVA. We believe ANOVA is less powerful than a paired t-test under these circumstances, particularly since the number of mice used both by us and by Dowd et al. was rather small. However regardless of the statistical test, the fact remains that the fold-differences in neutralizing titres were at best small (˜2-fold) for Dowd et al., but moderate (˜4-fold to ˜10-fold) for us, and the reasons for these differences are unknown.

In summary, our results suggest that the new American strains of Zika may relatively virulent, at least in juvenile mice. The American strain PRVABC59 is almost as virulent in mice as MR766, despite the fact that MR766 has been repeatedly passaged in mice, while PRVABC59 has not. Furthermore, we detect moderate serological differences between the ancestral African virus and the modern American Zika virus, with a four to ten-fold difference in neutralization titer when comparing homologous to heterologous viruses. On the one hand, a four to ten-fold difference is relatively modest. On the other hand, given that immunity can wane with time, and that individual responses vary, even a four-fold difference could be functionally important for a vaccine. In our studies, the number of mice used was small, and the serological results are based solely on the in vitro PRNT_50_ assay, so of course further and larger studies of various types are required. Nevertheless, since vaccines against Zika virus are already being used in human volunteers (NCT02840487, NCT02887482, NCT02809443), it is important to consider that for maximum effectiveness against the American virus, vaccines should be based on an immunogen from an American virus.

## Acknowledgments

This study was supported by a grant from the NIH (NIH 1R01AI110792 – 01A1) and also funded by the Center for Biotechnology, a New York State Center for Advanced Technology; New York State Department of Economic Development Contract #C150127; and corporate support.

